# Flexibility in movement strategies of neotropical nectarivorous birds: insights from high-Andean hummingbirds and flowerpiercers

**DOI:** 10.1101/2025.02.28.640730

**Authors:** Cristina Rueda-Uribe, Pedro A. Camargo-Martínez, Jonathan Espitia, Manuela Lozano-Rocha, Juan Pablo Ríos, María Ángela Echeverry-Galvis, Lesley T. Lancaster, Isabella Capellini, Justin M. J. Travis, Alejandro Rico-Guevara

## Abstract

Nectarivorous birds should have flexible movement behaviours in response to the presence of competitors and the spatiotemporal availability of flowering plants, particularly in tropical regions where flower blooms follow patterns of precipitation that are unpredictable across years. While pollinators such as hummingbirds (Trochilidae) have diet breadths that are constrained by trait-matching with flowers, nectar-robbing flowerpiercers are tanagers (Thraupidae) that typically drink nectar from holes they pierce near the flower’s base. Consequently, distinct movement patterns for these two bird families would be expected from optimal foraging theory, yet little is known about how tropical nectarivores move in response to fluctuating conditions. We used fine-resolution tracking data from an automated radio telemetry grid to compare movement patterns between hummingbirds and flowerpiercers in high-Andean mountain ecosystems. We obtained an accumulated total of 435,513 location estimates and 452 tracking days from 22 individuals across six different bird species. Our results indicate that hummingbirds exhibit a greater diversity of movement behaviours in comparison to flowerpiercers, with varying space use and recursion patterns that are characteristic of sedentary, commuting/traplining and exploratory strategies, whereas most species of flowerpiercers were classified as central-place foragers. However, daily movement metrics show that there is substantial variation, and hierarchical clustering does not necessarily group together bird families, species, nor even individuals as more similar to each other. Flexibility in daily movement behaviours has seldom been described for neotropical nectarivorous birds in the wild. It emerges as an important trait to adjust behaviour to variable local contexts, and may be adaptive for persistence in challenging mountain ecosystems where weather conditions are harsh and floral resources are seasonal and limited. A better understanding of flexibility in movement behaviour can enhance our predictions about how animals respond to environmental change and anthropogenic pressures.

## Introduction

Animals move to feed, find shelter, reproduce, avoid predation and competition, and respond to environmental conditions, but movement is energetically demanding and must be counterbalanced by the benefits gained (Nathan et al. 2008). Trade-offs between costs and benefits of movement have been studied under optimality approaches (Emlen 1966, MacArthur & Pianka 1966), within which optimal foraging theory has given a framework to understand how animals make decisions that maximise the energy gained from food and reduce the costs involved in movement, competition, and predation risks, ultimately maximising fitness (Pyke et al. 1977, Inouye 1978, Fontaine et al. 2008, King & Marshall 2022). In this sense, studying animal movement patterns not only provides ecological and evolutionary insights into species’ behaviours but can also be useful to understand whether animals may flexibly respond to change (Tucker et al. 2018). Despite the significance of studying animal movement, there are still significant knowledge gaps for certain regions and taxonomic groups (e.g. Hsiung et al. 2018, Jahn et al. 2020).

Nectarivorous birds are interesting to study in the context of optimal foraging (Pyke 1978, Pyke 2019) because they are endotherms with energetically demanding metabolisms fueled by nectar. Nectar is a replenishable yet ephemeral resource (given that plants periodically produce nectar, but nectar is also depleted by competitors), that follows temporal patterns of flower blooming and can be aggregated in space (Gill & Wolf 1979, Temeles et al. 2006, Schmid et al. 2016). Consequently, nectar-feeding birds should respond to spatiotemporal variation in nectar and local community contexts by adopting flexible movement behaviours. At shorter, daily time scales, behavioural flexibility can include adjusting time between flower visits to allow for nectar replenishment (Tello-Ramos et al. 2019), modifying duration and frequency of flower visits depending on nectar availability (Garrison & Gass 1999, Temeles et al. 2006), minimising distances in foraging routes (Tello-Ramos et al. 2015), and expanding foraging areas as a consequence of the presence of competitors (Hazlehurst & Karubian 2018). Seasonal flexibility in response to flowering blooms can occur by switching movement strategies across years (Smetzer et al. 2021) or travelling long distances to exploit seasonal resource availability (Guillamet et al. 2017).

Flexible movement behaviours of nectarivorous birds in response to flowering blooms should be particularly prevalent in the tropics, where floral phenology is shaped by precipitation regimes that are unpredictable across years (Stiles 1978, Sullivan et al. 2024) and the spatiotemporal patterns of blooming are highly variable among plant populations and species (Brown & Hopkins 1996, Sakai 2001). The complexity in spatiotemporal rhythms in tropical ecosystems offers interesting model systems to better understand behavioural flexibility in response to fluctuating resources, but movement strategies of tropical nectarivorous birds have only been thoroughly studied for just a few species.

Described movement behaviours of tropical nectar-feeding birds range from territorial sedentarism, with species remaining in a limited area which they usually defend (e.g. hummingbirds, *Aglaeactis cupripennis* (Shining Sunbeam), Hazlehurst & Karubian 2018, Céspedes et al. 2019), to central-place foraging, which refers to birds repeatedly returning to a central location, and commuting, with movement to feeding sites that are distinct from roosting or breeding areas (e.g. Hawaiian honeycreepers, *Himatione sanguinea* (Apapane) and *Drepanis coccinea* (Iiwi), Smetzer et al. 2021). In addition, traplining in hummingbirds is a strategy in which animals feed from flowers along repeatable routes, potentially covering long distances to visit flowers that they will typically not defend (Stiles 1975). Nomadism also emerges as a prevalent movement strategy to follow flowering blooms (Guillamet et al. 2017) and could potentially lead to explorative behaviours when they arrive in new areas. In fact, nectarivory has been identified as a factor that increases propensity of elevational movement in the Neotropics (Barçante et al. 2017), and there is accumulating evidence that even very small nectarivorous birds, such as hummingbirds, have widespread movements across ecosystems and elevations (Rueda-Uribe et al. 2023) and can achieve long-distance latitudinal and altitudinal migration (Williamson et al. 2024).

However, dietary specialisation limits an animal’s behavioural flexibility, for reliance on just a few types of food sources restricts the range of decisions it can make. In hummingbirds, the most speciose and oldest lineage of nectar-feeding birds (Hewes et al. 2022), trait-matching—both in length and curvature—largely determines from which plants the bird can most efficiently drink nectar from (Rico-Guevara et al. 2021), although both short and long-billed species also rob nectar from long flowers (Colwell et al. 2023, Reid et al. 2023). Analyses of hummingbird-plant interaction networks have shown that floral visitation by hummingbirds often correlates with increased matching between the lengths and shapes of birds’ bills and flower corollas (Maglianesi et al. 2014, Weinstein & Graham 2017), although bill morphology does not necessarily lead to resource specialisation because there may be multiple suitable resource species, imprecise ‘fit’ requirements, and context-dependent probabilities of encountering interaction partners (Dalsgaard et al. 2021, Hurtado et al. 2024). Nevertheless, hummingbirds have radiated across the Americas in close association with flowering plants (Barreto et al. 2024) and vary remarkably in the degree to which they have evolved specialised interactions with their plant partners, with some hummingbirds visiting only a few plant species and others having very wide diet breadths (Rodríguez-Flores et al. 2019, Leimberger et al. 2022).

Expected movement patterns that result from dietary specialisation are that generalist hummingbirds are able to feed from a smaller area which they will defend from intruders, while specialised hummingbirds fly over longer distances to visit select flowers along routes (Betts et al. 2015, Sargent et al. 2021). This distinction in hummingbird foraging strategies has led to the grouping of hummingbirds into territorialists and trapliners, respectively (Feinsinger & Colwell 1978), although this dichotomous classification is better viewed as a continuum given the high variation between and within species, and even within individuals, in response to contextual factors such as resource availability and competition (Sargent et al. 2021). In fact, hummingbird feeding preferences can be highly plastic and they often exhibit opportunistic behaviours (Abrahamczyk & Kessler 2015), including nectar-robbing by piercing flower bases or using holes previously made by other species (e.g. Pelayo et al. 2011, Reid et al. 2023). However, flexibility of hummingbird movement behaviours has been mostly studied at artificial feeders, with little to no information under wild conditions (but see Hazlehurst & Karubian 2018), which limits our understanding of how they respond to community and landscape contexts.

Even less is known about the foraging movement patterns of flowerpiercers, passerines of the genus *Diglossa* in the tanager family (Thraupidae) that pierce the base of flower corollas with their mandibles to drink nectar (Mauck & Burns 2009). Flowerpiercers compete with hummingbirds for nectar where their distributions overlap (Arizmendi 2001, Hazlehurst & Karubian 2018). As nectar-robbers, flowerpiercers may potentially overcome the limitations of trait-matching because they can extract nectar either from the front (similar to legitimate pollinators) or the side (by piercing holes) of flowers (Cuta-Pineda et al. 2021). Flowerpiercers as nectar-robbers may thus benefit from having broader diet breadths (Rojas-Nossa et al. 2016) which could allow for shorter foraging distances, thereby increasing foraging efficiency (Irwin et al. 2010). Still, research on flowerpiercer foraging strategies is notably lacking. It may be that flowerpiercers minimise the costs of movement by establishing smaller feeding areas like generalist hummingbirds, but there is evidence that flowerpiercers join mixed-species flocks (Hilty 2021, Vásquez-Ávila et al. 2021) and fly longer distances between habitat types (Rojas-Nossa 2007).

To address these knowledge gaps, our study used an automated radio telemetry system (ARTS) to collect fine-resolution movement data for multiple species of hummingbirds and flowerpiercers in natural landscapes. We aimed to investigate whether hummingbirds, as legitimate pollinators more constrained by their morphological match to flowers for nectar access, differ from nectar-robbing flowerpiercers, in terms of their daily movement patterns and individual- and species-level behavioural flexibility. For this, we conducted our study in a high-elevation mountain site in the tropical Andes, where flowering is seasonal (e.g. Gutiérrez Z. et al. 2004, Pelayo et al. 2021) and nectarivores ought to adjust to changing flower availability. We tagged three hummingbird species that differ in key functional traits, and reported foraging and movement behaviours: *Chalcostigma heteropogon* (Bronze-tailed Thornbill) is a short-billed and medium sized hummingbird that can hold territories but also trap-lines visiting scattered flowering plants (Hilty & Brown 1986, Hilty 2021), and commonly clings to flowers or walks on the ground to feed instead of hovering (Colwell et al. 2023); *Colibri coruscans* (Sparkling Violetear) has a medium body mass, medium-length bill, and it is notably aggressive; it defends territories by perching on high branches and vocalising frequently (Hainsworth 1977, Hilty 2021); and *Pterophanes cyanopterus* (Great Sapphirewing) a large, long-billed hummingbird that holds territories when there are high nectar rewards (Snow 1983, Gutiérrez Z. et al. 2004) but has also been reported to trap-line, and glide along mountain slopes, potentially to reach distant areas (Stiles 2008, Hilty 2021). In addition, we tagged three flowerpiercer species: *Diglossa cyanea* (Masked Flowerpiercer), which has the least hooked bill of the three flowerpiercer species we tracked, and is reported to be more frugivorous, join mixed species flocks and does not hold territories (Rojas-Nossa 2007, Hilty 2021); *Diglossa humeralis* (Black Flowerpiercer), the smallest of the three flowerpiercer species and with the thinnest bill (relative to head size, Hilty 2021), is not reported to often join mixed species flocks and is territorial when single (non-breeding) and in pairs (when breeding, Hilty & Brown 1986); and *Diglossa lafresnayii* (Glossy Flowerpiercer), which seems to be territorial when single or in pairs but can also join mixed species flocks (Hilty 2021). Although flowerpiercers are morphologically more homogeneous than hummingbirds, their bill length and hook shape is related to the frequency with which they extract nectar by piercing holes in the sides of corollas or by inserting their bills in the flowers from the front (Rojas-Nossa 2007), yet more research is needed to establish if these behavioural differences are also related to diet breadths and specialisation, and not only feeding efficiency (Schondube & Martínez del Rio 2003).

If hummingbirds are more limited in the variety of flowers they can feed from in comparison to flowerpiercers, we predicted that hummingbirds would be forced to cover greater areas and travel longer distances during the day, selecting areas of elfin forest, which probably are higher quality resource patches because they have a greater diversity of ornithophilous flowers compared to the paramo in this site (Rueda-Uribe et al. in review). Also, the need to follow the spatiotemporal arrangement of their preferred flowers would result in hummingbirds holding looser territories that could change location across days and in which they would spend less time and return to fewer times and at more prolonged intervals. Given that we tracked three different species of hummingbirds that differ in key functional traits, we expected high variation in movement strategies across hummingbird species but not flowerpiercers, because the latter are presumably more similar in their morphological adaptations, and thus would also have similar foraging behaviours and related movement strategies. However, given that *D. humeralis* has been previously reported to not join mixed species flocks as frequently as the other two flowerpiercers, we predicted that *D. humeralis* would be more sedentary. Finally, we predicted that all individuals should exhibit some extent of flexibility in their daily movement patterns in response to the spatiotemporal variability of nectar availability in high-elevation mountainous ecosystems.

## Methods

### Bird tracking

Birds were tracked with an ARTS grid installed in the Valle de los Frailejones, located at 3,162– 3,391 m a.s.l. inside Chingaza National Natural Park in the Eastern Cordillera of Colombia (4°31′30.5′′N 73°46′23.9′′W), and contains high-Andean ecosystems of paramo and elfin forest. The ARTS grid covers an area of 0.72 km^2^, with 46 receiving nodes (Node V2, Cellular Tracking Technologies - CTT, USA) set approximately at 2.5 m above the ground and 150 m apart (Figure 1A). These nodes continuously detect individual tag ID signals and radio signal strengths (RSS, measured in decibels) at 433 MHz and a maximum distance of 300 m, making the area of coverage of the grid approximately 2.05 km^2^, albeit limited by interference from topography and vegetation and increased localisation error outside of the grid (Rueda-Uribe et al. 2024). Nodes communicate with a base station (CTT SensorStation) that has four receiving Yagi antennas (∼10 m above the ground), from which data is downloaded manually.

**Figure 1.**
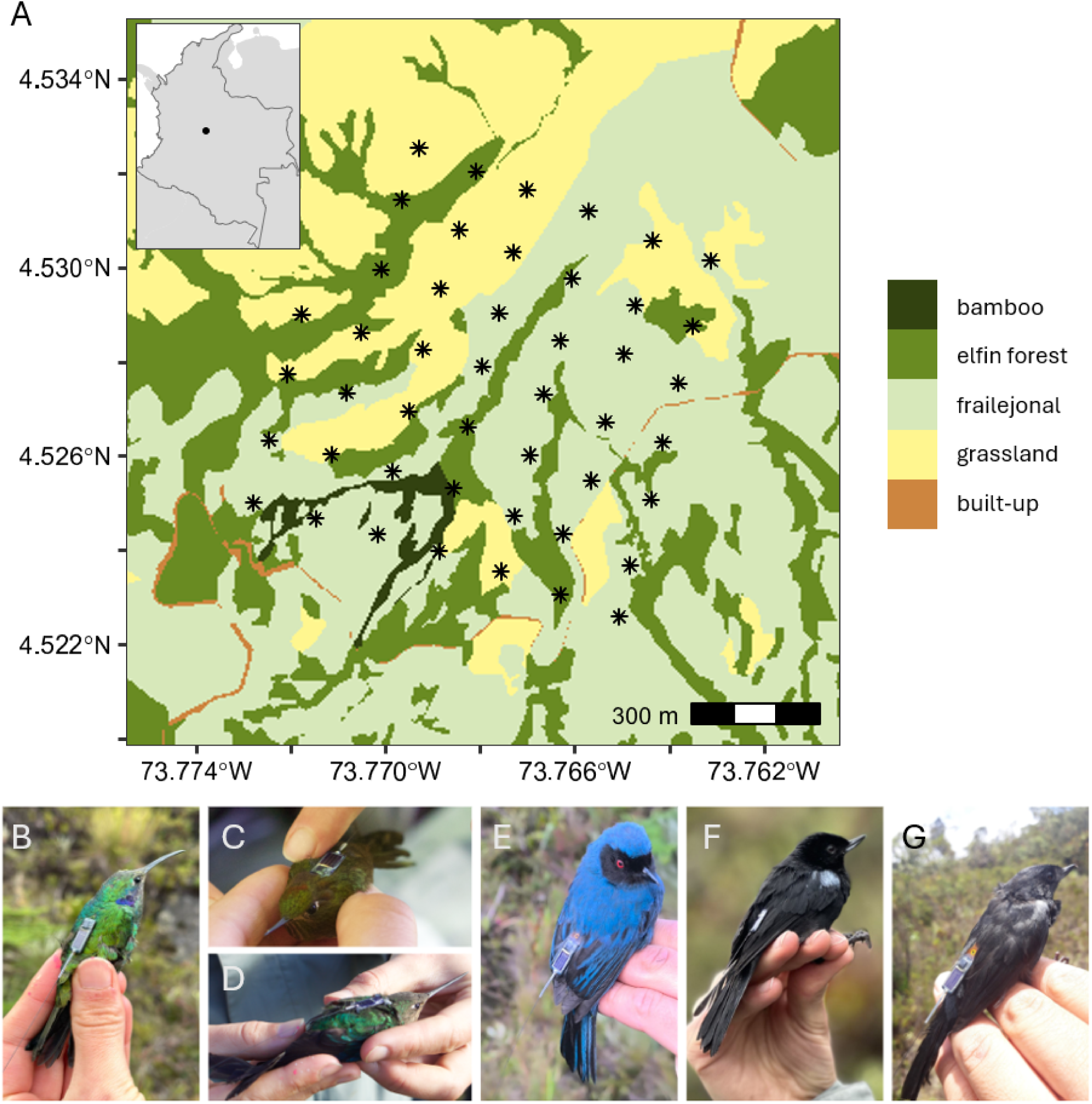
Study site and species of tagged nectarivorous birds. A) Location of automated radio telemetry grid in the Valle de los Frailejones, Chingaza National Natural Park in Colombia. Receiving nodes that compose the grid are shown as asterisks, and vegetation covers of the valley are mapped at a 5 x 5 m resolution and coloured according to the legend. B-G) Tagged hummingbirds: B) *Colibri coruscans* (Sparkling Violetear), C) *Chalcostigma heteropogon* (Bronze-tailed Thornbill), and D) *Pterophanes cyanopterus* (Great Sapphirewing). Flowerpiercers: E) *Diglossa cyanea* (Masked Flowerpiercer), F) *Diglossa lafresnayii* (Glossy Flowerpiercer), and G) *Diglossa humeralis* (Black Flowerpiercer).

Two different kinds of tags were used to track birds, LifeTags and PowerTags (CTT). LifeTags rely on a miniaturised solar panel to emit a radio signal every 2-5 s and have a mass of 0.35-0.45 g including the harness. PowerTags use a battery, were set to emit a signal every 60 s, and have a mass of 0.35 g. Tags were attached to birds captured with mist nets (32 mm mesh, 2.5 × 6, 9, or 12 m). Mist nets were open in the mornings (6-11 am) and checked every 15-20 min, during different field seasons between December 2022 and August 2024 (Table S1). All birds were handled by trained experts with research and ethics permits (see Ethics Statement).

We tagged 24 adult birds from three different hummingbird species and three flowerpiercer species (Figure 1B-G): six *Ch. heteropogon*, five *Co. coruscans*, five *P. cyanopterus*, two *D. cyanea*, four *D. humeralis* and two *D. lafresnayii* (Table S1). Tags were attached to flowerpiercers using a leg loop harness with 0.6 mm cotton stretchy string (Naef-Danzer 2007). For *P. cyanopterus*, tags were attached with the chest and wing harness described in Williamson and Witt (2021), with a 3D printed chassis designed by Rueda-Uribe et al. (2024) and 0.5-0.7 mm StretchMagic string. For *Co. coruscans* and *Ch. chalcostigma*, the tags were glued on the scapular area of their backs with non-irritating superglue (Loctite PureGel, Hadley & Betts 2009). Deployed tags were approximately within 3%-5% of bird body mass, and appropriate tag fit was always checked. These types of tags have been shown to not significantly affect flying, feeding, preening, and perching time budgets in a species of hummingbird (Sargent et al. 2025). Hummingbirds were fed sugar water before release and all birds flew normally after being released.

### Track estimation and movement metrics

Maximum RSS calculated over 10 s for LifeTags and 120 s for PowerTags was used to estimate the location coordinates of individual animals, based on a previously calibrated signal-distance decay relationship and using a smoothing and multilateration workflow described in Rueda-Uribe et al. (2024). Resulting tracks were further cleaned by using a 25 m/s speed threshold to remove outliers. This threshold was chosen because 95% of observed speeds were under this value, and it is a feasible natural limit of bird flight speed for these species (see Rueda-Uribe et al. 2024). Moreover, tracks from PowerTags were filtered to exclude relocations during the nighttime so that all analyses were performed for daytime behaviour. We removed PowerTag relocations that were before the minimum (5:44) and after the maximum (18:09) times of day recorded with LifeTags. Additionally, birds that were tracked for fewer than two days and days with less than one hour of accumulated tracking time were deemed as not representative of daily movement behaviours, so they were excluded from further analyses (Table S1).

Seven movement metrics were calculated to describe daily movement behaviours. Space use metrics included the daily area utilised by individuals (1), cumulative distances travelled during the day (2) and displacement of daily areas in the landscape throughout the tracking period (3). Daily utilisation distributions were calculated with 95% autocorrelated kernel density estimates (AKDEs) using the package “ctmm” in R (Fleming & Calabrese 2023). AKDEs account for autocorrelation in movement tracks, which is especially important for data that is collected at high sampling frequencies such as that from an ARTS grid. Also, AKDEs can incorporate error in track estimation, be used to compare individuals that have different sample sizes and sampling rates, and provide 95% confidence intervals to report uncertainty in downstream analyses (Fleming et al. 2015, Silva et al. 2022). AKDEs were not calculated when continuous-time movement models failed to converge (Silva et al. 2022) or if birds had fewer than 15 relocations for that day. In addition, resulting AKDEs were manually checked by visually inspecting them on a map with the raw location points, so that AKDEs that covered areas with no points or low densities of points were considered to be overestimating utilisation distributions and were thus filtered out.

Daily area was measured as the area covered by 95% AKDEs. Daily cumulative distance was the summation of distances between track relocations, and was standardised by tracking duration because these two variables were correlated, with cumulative distance increasing the longer a bird was tracked (Figure S1). Displacement of daily areas in the landscape was measured as the distance between each centroid of daily 95% AKDE polygons and the centroid of aggregated areas used throughout the tracking period. Recursion metrics were calculated by defining daily core use areas as 50% AKDEs, and estimating the residence time (4) as the proportion of time a bird spent inside its core area, revisitation rate (5) by counting how many times it exited its core area and returned divided by track duration, and mean return time (6) as the average duration of forays outside its core area.

Habitat use was measured as the proportion of elfin forest within individual daily core use areas (7). Forest was selected as the most suitable vegetation cover to include in movement metrics as habitat use because in this valley it contains a higher diversity and abundance of flowers that provide nectar to birds (Rueda-Uribe et al. in review), and none of the abundant hummingbird-pollinated plants of other vegetation covers (*Espeletia grandiflora* in the “frailejonales” cover and *Puya* sp. in grasslands) were blooming during the tracking period. Forest proportion was calculated as the amount of this vegetation cover in square metres divided by total area. Forest cover was extracted from mapped vegetation covers for the area within and surrounding the ARTS grid. We generated the vegetation map by drawing polygons over Google Earth Pro satellite imagery that identified vegetation of grassland, bamboo, “frailejonales” (rosetted paramo plants), elfin forest trees and shrubs, and “built-up areas” (a small dirt road and park ranger housing). Polygons were transformed into a 5 x 5 m raster covering a buffer area of 300 m around the grid, using spatial analyses packages “terra” (Hijmans 2024) and “sf” (Pebesma & Bivand 2023) in R.

### Variation in movement behaviour

To test for differences in movement behaviours between hummingbirds and flowerpiercers, we fit linear mixed effects models for each of the seven daily movement metrics explained above as response variables, with the grouping of birds by family as fixed effects. Response variables were transformed by the square root or log10 to approach normality. Individuals nested within species were included as random effects on the intercept. We calculated 95% confidence intervals to check if estimated effects differed from zero, tested the models against a null (model without the fixed effect of bird family) with an Analysis of Variance (ANOVA) test and ranked them according to likelihood and Akaike Information Criterion (AIC). The models were fitted with the package “lme4” in R (Bates et al. 2015) and temporal autocorrelation was checked by plotting autocorrelation in residuals.

Additionally, we used agglomerative hierarchical clustering and Principal Components Analysis (PCA) of the seven movement metrics to identify variation in movement strategies (Abrahms et al. 2017, Smetzer et al. 2021). For this, we first standardised the metrics by scaling around their mean. Then, we clustered these scaled values with the function “hclust” in R, setting the algorithm method to “ward.d2”, which uses Ward’s minimum-variance criterion with squared clustered dissimilarities (Murtagh & Legendre 2014). We assessed the support of each cluster by calculating p-values with 1,000 multiscale bootstrap resampling with the package “pvclust” in R (Suzuki et al. 2019), setting *a priori* that clusters had sufficient support if their p-value was lower than 0.05 for 90% of iterations. Finally, we visualised variation by calculating an Euclidean distance matrix to perform a PCA on the seven movement metrics with the function “prcomp” in R and selected the two components that together explained the highest proportion of observed variation. By plotting variable loadings and observed values in a two-dimensional plot, we visualised the location of observations and calculated convex hulls as proxy to assess functional space areas occupied by individuals, species and nectarivore groups. Hierarchical clustering and PCA were performed separately on daily metrics (several values per metric per individual) and mean values by individual (one value per metric per individual).

## Results

We estimated tracks for 22 birds of three hummingbird and three flowerpiercer species, for a total of 435,513 location estimates after track filtering that summed up to 452 days of tracking across individuals. The amount of location estimates and the duration of tracking varied for individual birds, with some having tracking data only for a couple of days while others being tracked for over 3 months, with the maximum being 97 days from tag attachment. Data gaps of some days and subsequent reappearances suggest that individuals leave the area and later return. Moreover, very low signals or detections from just a few nodes located in the edges of the grid are most probably the result of individuals being present in the broader area but not inside or near the grid (Figure S2). For two birds, tags did not generate maximum RSS reads strong enough to estimate location information (minimum node-specific values range between −100 and −110 dB for distance estimation and subsequent multilateration, see Rueda-Uribe et al. 2024). From the resulting tracks, 306 daily utilisation distributions were calculated with AKDEs for 14 individuals with more than two days of tracking, and this was the data used in downstream analysis (Table S2).

Our analysis of daily movement behaviours showed that, on average, hummingbirds covered greater daily areas compared to flowerpiercers (0.73 ± 0.66 vs. 0.48 ± 0.28 km^2^) but travelled shorter cumulative distances (3.35 ± 2.94 vs. 6.19 ± 3.93 km/h) and had very similar centroid displacements across the landscape (0.19 ± 0.10 vs. 0.17 ± 0.03 km). In terms of recursion metrics to core areas (50% AKDEs), hummingbirds on average spent more time in core areas (0.60 ± 0.20 vs. 0.51 ± 0.10) but also took longer to return after leaving their core areas (17.66 ± 10.70 vs. 8.67 ± 12.10 min), and had lower revisitation rates (3.00 ± 3.08 vs. 7.07 ± 5.90 revisits/h) compared to flowerpiercers. Both groups had similar proportions of forest cover inside their core areas (0.30 ± 0.11 vs. 0.28 ± 0.05; Figure 2). However, linear mixed effects model results comparing space use metrics between the two bird groups were not different from the null and did not have effects that were different from zero with 95% confidence intervals (Table S3). Variation among individuals was greater than across species for most metrics, with individual and species accounting for 8-54% and 0-41% of variance in random effects, respectively (Table S4, Figure S3).

**Figure 2.**
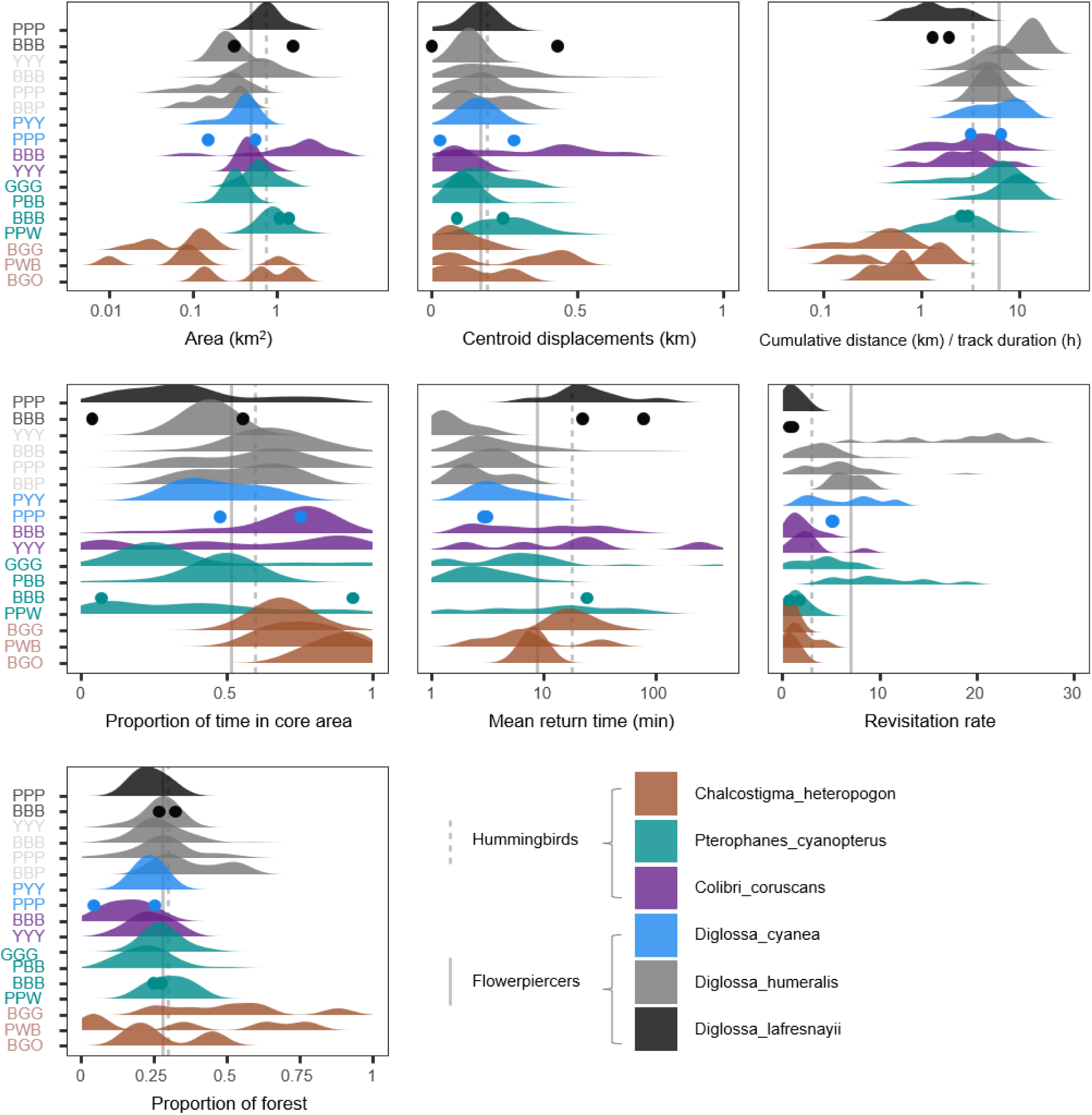
Distribution of seven daily movement metrics of tracked individual hummingbirds and flowerpiercers. Each individual has a three-letter code and is represented in the y axis, measured movement metrics are on the x axis. Note that the metrics of area, cumulative distance, and mean return time have a logarithmic scale to aid visualisation. Species are coloured according to the legend, and vertical lines show mean values for hummingbirds (dashed lines) and flowerpiercers (solid lines). Three individuals had information for only two days and thus their space use metrics are represented by points rather than distribution curves. They are included here for visualisation but excluded from analyses.

Clustering average movement metrics by individual resulted in three clusters that were strongly supported by multiscale bootstrap resampling probabilities (90, 92 and 95% of iterations with p < 0.05, Figure 3A). However, clusters grouped together different bird groups and species, with only one cluster containing a single species. This cluster grouped together all *Ch. heteropogon*, which showed typical sedentary behaviour, spending longer times in core areas and having lower daily travel distances than the other clusters (Figures 3A and 4A). Most flowerpiercers (except *D. lafresnayii*), together with one *P. cyanopterus*, were grouped in a cluster characterised by behaviours related to central-place foraging, with short, frequent revisits to a central core area, and higher daily distances travelled. The third cluster, which grouped two *P. cyanopterus*, one *Co. coruscans* and one *D. lafresnayii*, can be described to have a commuting or traplining strategy, since they covered greater areas, had longer return times, short residence times, and lower revisitation rates to core areas, while travelling shorter cumulative distances in comparison to central-place foragers (Figure 5). The only bird that was not clustered with other individuals was one *Co. coruscans*, which covered very large daily areas and had long distances between daily centroids. Clustering of daily measures by individual resulted in 64 clusters (equal or greater than 95% iterations with p < 0.05), with all individuals grouped in more than one cluster (Figure 3B).

**Figure 3.**
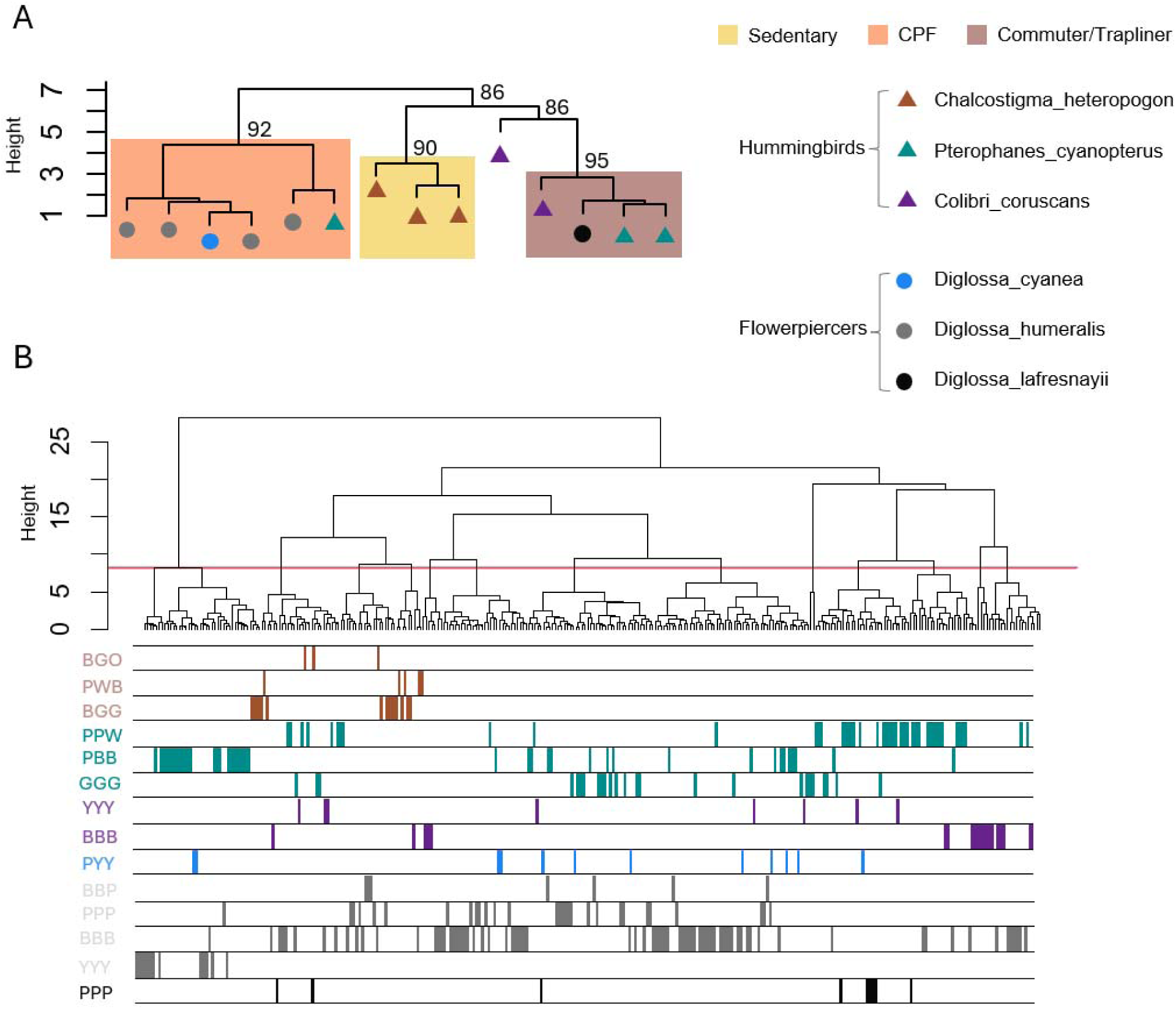
Hierarchical clustering dendrogram of seven daily movement metrics of tracked hummingbirds and flowerpiercers. A) Clustering with mean values by individual. Each point represents an individual, and species are coloured according to the legend. Identified clusters are shown as rectangles and labelled according to movement strategies of central-place foraging (CPF), sedentarism and commuting/traplining that were described with principal components analysis (see Figures 4 and 5). Numbers show cluster support as the percentage of iterations with p > 0.05 in multiscale bootstrap resampling with 1,000 iterations. B) Clustering with raw values, with several values per individual. Sixty-four clusters were identified, all of which were under a maximum height of 8.18, shown as a horizontal red line. Bars under dendrogram show presence of individuals across clusters, with each horizontal division separating individuals identified with a three-letter code and the colours indicating species according to the legend.

**Figure 4.**
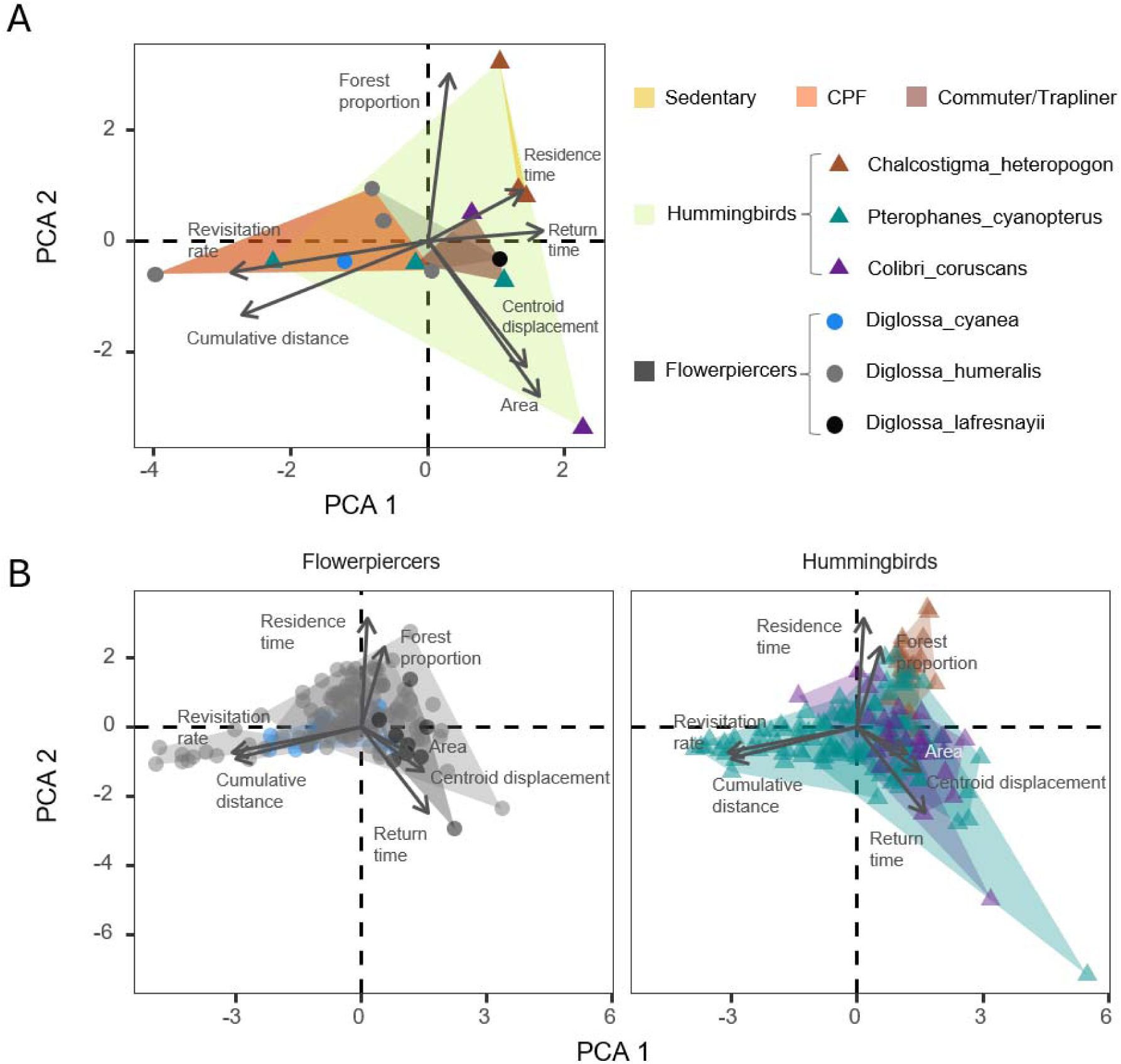
Principal components analyses (PCA) and hierarchical clustering of seven daily movement metrics of tracked hummingbirds and flowerpiercers. A) Analyses based on mean values by individual, with each point representing an individual, coloured according to species and family as shown in the legend. Functional space occupied by bird family is shown as a convex hull coloured in light green for hummingbirds, grey for flowerpiercers, and three identified groups from hierarchical clustering in yellow (sedentary), orange (CPF = central-place forager) and dark red (commuter/trapliner). B) Analyses including all values by individual with points showing daily values, and separated in different panels by flowerpiercers and hummingbirds to ease visualisation. Refer to Figure S4 for panels separated by species, and Figures S5-S7 for plotting of a third PCA component. The functional space occupied by an individual is shown as a convex hull and coloured according to species, as shown in the legend. Note that variable loadings were multiplied by 5 to improve visualisation (raw values are available in Table S5).

**Figure 5.**
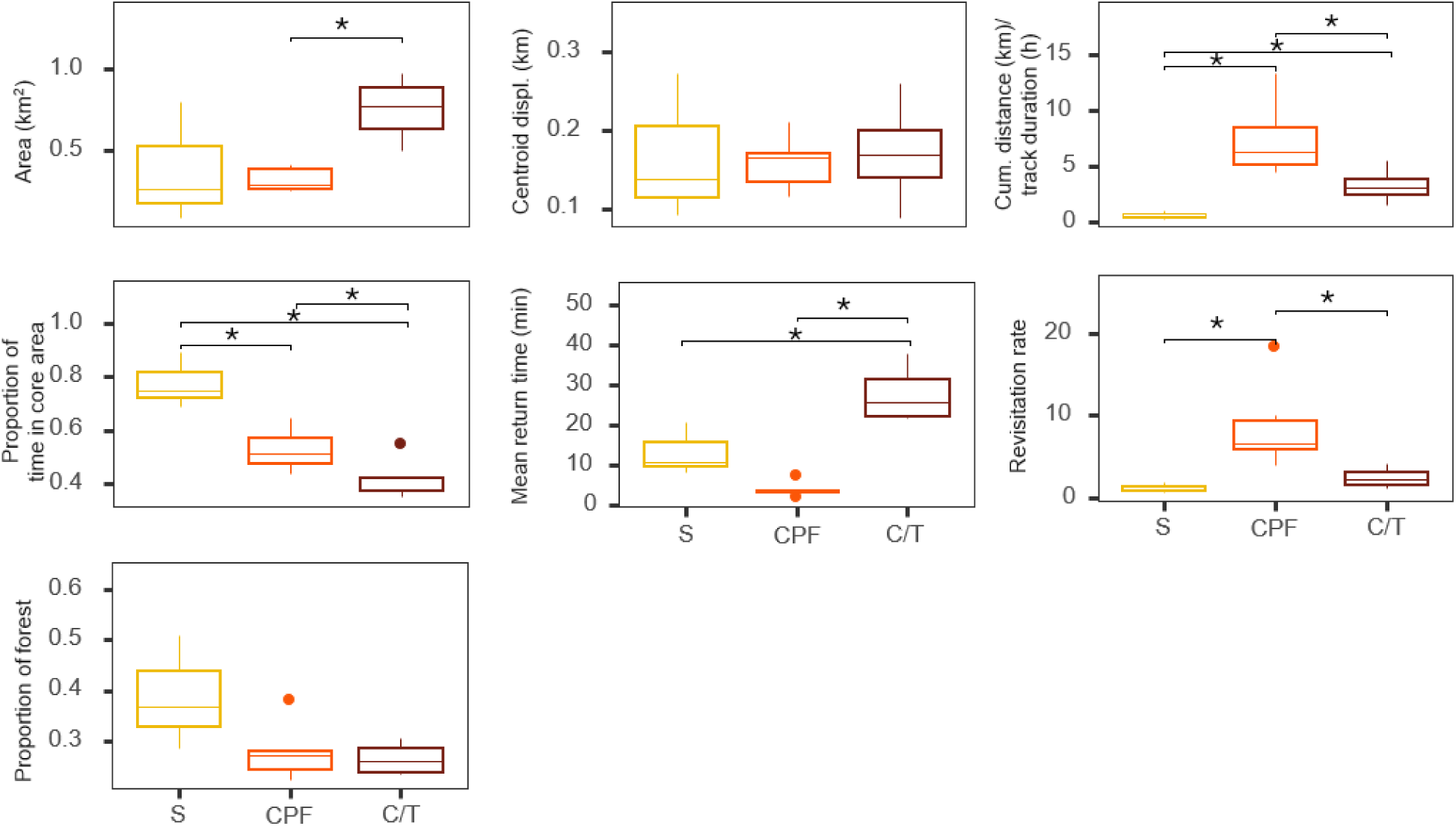
Average values of seven daily movement metrics according to movement strategy groups of tracked hummingbirds and flowerpiercers. Boxplots show first and third quartiles with lower and upper box hinges, median values with middle horizontal lines, 1.5 interquartile ranges with vertical lines, and outliers with points. Asterisks show significant differences in the mean between groups, with p < 0.05. Refer to Figures 3 and 4 for species identified within each group by hierarchical clustering. S = Sedentary, CPF = Central-Place Forager, C/T = Commuter/Trapliner.

Variance among daily movement patterns was explained up to 69 and 51% by the first two principal components of the PCA for the mean and daily movement metrics, respectively. Plotting hummingbirds and flowerpiercers in the PCA plot shows that hummingbirds occupy a greater functional space in terms of their movement metrics compared to flowerpiercers, measured as the area of convex hulls for each bird group (16.22 vs. 4.75, Figure 4A). However, there is overlap across all individuals (Figure 4B), and *Ch. heteropogon* has the lowest variability in observed movement metrics (Figure S4).

## Discussion

Our fine-resolution tracking enabled the comparison of daily movement behaviours for the two main groups of neotropical nectarivorous birds, both of which have been largely understudied in their movement ecology. Using seven different metrics to describe movement, we found that hummingbirds exhibit greater interspecific variation in space use, recursion behaviours and habitat associations in comparison to flowerpiercers, probably as a result of morphological and ecological specialisation. However, hummingbirds as a group did not significantly differ from flowerpiecers in any of the analysed daily movement metrics, in part because some species of hummingbirds overlap with flowerpiercers in their movement strategies, but also due to substantial variation within individuals and species for both bird families. Importantly, our findings reveal flexible movement behaviours that have seldom been described for nectarivorous birds in the wild.

The observed movement behaviours in our study support previous evidence that hummingbirds have a diversity of foraging strategies spread along a continuum (reviewed in Sargent et al. 2021). The three hummingbird species that we tracked notably vary in bill lengths, body mass and wing chord (Figure S8), of which bill length, in particular, is a key functional trait in determining which plants they can feed from (Rico-Guevara et al. 2021, Dalsgaard et al. 2021). We found that *Ch. heteropogon*, the short-billed and smaller species, exhibited characteristically sedentary behaviour by moving over smaller areas, travelling shorter distances, and remaining within core areas. This species has been previously reported to visit a broad diversity of plants in this study region in comparison to the two other hummingbird species we tracked (Manrique-Garzón et al. 2025), supporting our hypothesis that generalist hummingbirds can reduce the costs of movement by foraging over smaller areas.

In contrast, we found that *P. cyanopterus* and *Co. coruscans* have movement behaviours that predominantly range between central-place foraging, commuting/traplining and explorative in our study landscape. Note that with our selected movement metrics, traplining, which is believed to be a common foraging strategy in hummingbirds (Sargent et al. 2021), cannot be distinguished from commuting, given that for both behaviours we would expect birds to spend most time outside of core areas and cover larger daily areas while travelling shorter cumulative distances in comparison to central-place foragers. Analysing movement behaviour at a finer temporal scale, such as by calculating travel speeds and space use within days, could be used to differentiate traplining from commuting behaviours. With our approach, we found that individuals of *P. cyanopterus* and *Co. coruscans*, in addition to their predominant movement behaviours, exhibited high variation in movement metrics across time and there was also evidence of sedentary behaviours for some days. The seasonal appearance of high-reward flowers can play a role in individuals temporarily remaining in a smaller area (Stiles 1985), as can be the case with the large inflorescences of bromeliads in the genus *Puya*, which *P. cyanopterus* frequently visits (Restrepo-Chica & Bonilla-Gómez 2017, Manrique-Garzón et al. 2025) or the abundant blooms of the melastome *Chaetogastra grossa*, which we have also observed to be notably associated with the presence of this hummingbird in the area. Inflorescences of *Puya goudotiana* were available when the *P. cyanopterus* classified as a central-place forager was tracked (June-July 2023), and the other two individuals of this species that were better described as commuters/trapliners generated tracking data while in our study landscape *C. grossa* was in bloom (December 2022-February 2023). Consequently, these interspecific differences in movement strategies for *P. cyanopterus* may be seasonal and respond to the spatial distribution of their floral resource, for plants of *Puya* sp. tend to be spatially aggregated (Restrepo-Chica & Bonilla-Gómez 2017), and could thus be exploited by hummingbirds remaining in their vicinity and exhibiting short, frequent revisits to a central core area. Movement behaviours of *P. cyanopterus* potentially also include nomadic movements to other areas (e.g., at lower elevations; Gutiérrez-Zamora 2008), which would explain several tagged birds leaving the area (Figure S2), but longer-distance movements for this species still need to be described.

For *Co. coruscans*, one individual remarkably covered very large daily areas, separated by long distances in the landscape. Given that this species has a seasonal occurrence in our study site, with individuals arriving around August and virtually disappearing in January (Rueda-Uribe et al. in review), it might be that this particular individual had recently arrived and was exploring. Although largely understudied, there are reports of *Co. coruscans* moving towards high-elevation ecosystems such as the paramo in the second half of the year (Gutiérrez-Zamora 2008). *Co. coruscans* is commonly described as a territorial species (Sargent et al. 2021), but our data shows that individuals within this species are not consistently sedentary across days and this is not necessarily the dominant behaviour during the period birds were tracked, even though they are very vocal in our study site, and we have seen them doing courtship displays. Future research could determine if this difference is due to exacerbated dominance and territorialism in rural and urban settings (Maruyama et al. 2019), where their behaviour is most frequently studied (e.g., Tellería & Garitano-Zavala 2024), or whether this variation in movement strategies is related to their breeding seasons.

Movement behaviour changes throughout an individual’s annual cycle, albeit in different ways for males and females in the absence of biparental care. In hummingbirds, it is mostly only the female that tends to the nest (Snow 1974, Baltosser 1986), while in flowerpiercers both sexes participate in nest defence and feeding young, although it seems that only females incubate (Schondube et al. 2003). In our study, all tracked individuals were adults and most were males, although there were also some individuals with unknown sex (Table S1), especially in the case of flowerpiercers since the studied species do not have sexually dimorphic plumage. Similarly, the only birds we tagged that exhibited signs of being in their breeding season were both individuals of *D. cyanea*, one had a visible cloacal protuberance and the other a refeathering brood-patch (scale 5, Deeming & Du Feu 2008). In the future, differentiating behaviours between sexes, for example by determining sex genetically, can be useful to identify sex-related differences in movement behaviours that may be related to breeding cycles, or other life histories stages.

In contrast to hummingbirds, most flowerpiercers were grouped in a cluster characterised by higher revisitation rates to core areas and cumulative distance travelled during the day, which is typical of central-place foragers (Stephens & Krebs 1986). We had predicted that the broader diet breadths of flowerpiercers would lead to sedentary behaviours because they can minimise the costs of movement by covering smaller areas. The advantage of central-place foraging over sedentarism, despite the increased energy expenditure of travelling longer cumulative distances throughout the day, may include decreasing the costs of defending a territory (Paton & Carpenter 1984) and gaining benefits from finding flower patches with greater rewards in different areas, while tending to a core area that individuals are familiar with (Smetzer et al. 2021) and may contain a nest. The single flowerpiercer that was not grouped with the rest was *D. lafresnayii*, due to this individual exhibiting longer return times to core areas and travelling shorter daily distances despite covering greater areas. This species, together with *D. cyanea*, is somewhat larger (>15g, Figure S8) and has been reported to join mixed species flocks, which would agree with our classification of this species in the commuting/traplining cluster. In the Andes, several species of birds join mixed species flocks that are generally unstructured and move continuously while foraging (Arbeláez-Cortés et al. 2011, Montaño-Centellas et al. 2023). There are several advantages of joining mixed species flocks, with the major benefits in terms of energy uptake being visiting different resource patches with increased foraging efficiency and lower predation risk (Mangini et al. 2023), but also possible costs such as competition with other members of the flock (Colorado & Rodewald 2014).

For both groups of nectarivorous birds, our findings show that there is high variation within species, and that individuals are flexible in their daily movement strategies, which most probably is an advantage if their main food resource depends on flowering rhythms and is patchily distributed in space. Particularly in the tropics, flexibility in movement behaviour for nectarivorous birds has been proposed as a mechanism for dealing with the low predictability of flowering across years (Smetzer et al. 2021). Although there is some evidence that hummingbirds adequate foraging behaviour to resource density (Wolf et al. 1976, Stiles 1985), flexible movement strategies remain poorly described for nectarivorous birds in general, being thoroughly described only for Hawaiian honeycreepers. This group of tropical nectarivorous birds has been shown to have seasonal associations to flowering plants (van Dyk et al. 2019), exploit flowering phenologies across elevations (Guillamet et al. 2017, Paxton et al. 2020), and switch between sedentary, central-place foraging and commuting strategies across years (Smetzer et al. 2021).

By measuring movement behaviours with a daily temporal resolution, our study revealed within-season flexibility, with birds adopting different strategies in a short period of time and even switching from one day to another. This suggests that birds are not only responding to seasonal variation in flowering phenology but also to local conditions that change at finer temporal scales, such as weather and the presence of competitors. Weather conditions in high-mountain tropical ecosystems are harsh and notoriously variable during the day, with strong wind gusts, lower air density, changes in cloud cover and rainfall, and temperatures that fluctuate from below freezing to over 20 °C in a matter of hours (Buytaert et al. 2006, Sklenář et al. 2015). All these abiotic factors affect bird flight efficiency, energy metabolism, levels of activity, and morphology (Altshuler et al. 2004, Groom et al. 2017, Stiles 2008, Beltrán et al. 2022). On the other hand, both exploitative and interference competition can elicit immediate changes in movement behaviour given the spectrum of floral foraging strategies (Rico-Guevara et al. 2021, Sargent et al. 2021). Nectarivorous birds can search for nectar over larger areas when resource availability is reduced by exploitative competition (Hazlehurst & Karubian 2018), change the duration and frequency of flower visits depending on the presence of competitors (Temeles et al. 2006), engage in aggressive behaviour (Wolf et al. 1976), and/or move elsewhere to avoid territorial disputes (Mac Nally & Timewell 2005). Competition for nectar in high-Andean ecosystems has been described as a main driver of hummingbird community assemblages (Lessard et al. 2016, Guevara et al. 2023) and is probably intense, for the abundance and diversity of floral resources decreases with elevation (Cuesta et al. 2017). Yet there are still several species of hummingbirds and flowerpiercers exploiting flower nectar in high-mountain ecosystems, potentially co-occurring through narrow niche partitioning (Montenegro Muñoz et al. 2015, Barros et al. 2020). We have recorded 12 species of hummingbirds and four flowerpiercers in our study site during 2022-2024 (Table S6). In addition, both groups of birds supplement their nectar-based diets with arthropods, and flowerpiercers also eat fruit (Stiles 1995, Schondube et al. 2003, Rico-Guevara 2008). This means that both hummingbirds and flowerpiercers, despite relying mostly on nectar for energy, must also be affected by the spatiotemporal changes in abundance of fruits and arthropods (e.g., Maya-García et al. 2024). Taken together, flexibility to constantly fluctuating conditions in high tropical mountains is probably essential for nectarivorous birds to survive, but movement behaviours have so far been vastly understudied in these ecosystems.

With this study we described daily movement behaviours for six species of neotropical nectarivorous birds that were previously unknown, and we identified substantial flexibility in movement behaviours within individuals. Our research can be further improved by extending the duration of tracking to monitor movement behaviour across years, while tracking more individuals and species. Limited sample sizes are a common challenge in tracking studies due to the loss of transmitters, technical failures, and, in the case of ARTS grids, individuals leaving the study landscape. However, GPS tracking is not yet possible with most hummingbird species and even some flowerpiercers due to their small body mass (but see Williamson et al. 2024), so ARTS grids are a practical option for fine-scale tracking of small nectarivorous birds (Smetzer et al. 2021), despite their limitations. In the future, the systematic use of our ARTS grid will generate more data across individuals and species, and also allow us to identify abiotic and biotic drivers of movement. Our effort to tag these high-Andean species, several of which have small ranges and most probably have never been tracked before, is an important step towards a better understanding of movement ecology in these valuable yet understudied ecosystems. Movement data gives information on how animals fulfill ecological needs in heterogeneous environments (Paxton et al. 2023), and can be used to better understand how animals will respond if the availability and abundance of resources is altered by anthropogenic change.

## Acknowledgements

We are grateful to all students, park rangers, and volunteers that supported and participated in fieldwork, particularly Juan Camilo Bonilla, Sarah Chaves, Angie Rodríguez, Alyssa Sargent, Aaron Skinner, Ana Melisa Fernandes, Miguel Ángel Muñoz-Amaya, Daniel Mancera, Fredy García, Luisa Díaz, Jessie Williamson, Rebekka Allgayer, Thomas Fey, Adriana Rueda, and Laura Manrique. We also thank Chingaza Parque Nacional Natural for all the logistical support and access to fieldwork permits, and Premium 3D for designing a tag chassis specifically suited for hummingbirds. NERC, BBSRC, Rufford Foundation, University of Washington Department of Biology, Washington Research Foundation, and School of Biological Sciences University of Aberdeen funded this work.

## Ethics statement

All birds captured with mist nets were handled by trained experts with research and ethics permits (Pontificia Universidad Javeriana FUA #127-22, University of Washington IACUC protocol number: 4498-05, University of Aberdeen approval 01/19/22). Research in Chingaza National Natural Park was carried out under the permits #20212000005863 and #20237160002153.

## Author contributions

C.R.U., P.A.C.M., M.A.E.G, J.M.J.T, and A.R.G. conceived this study, C.R.U., P.A.C.M., J.E., M.L., J.P.R. collected the data, C.R.U. designed the statistical analysis, analysed the data, created figures and wrote the original manuscript. M.A.E.G, I.C., L.T.L, J.M.J.T., and A.R.G. supervised the research. All authors revised and edited the manuscript.

## Data availability

All data and code will be made available in the Dryad Digital Repository and Zenodo.

## Collaboration ethics statement

This work is the result of a diverse collaboration that included people from different career stages, genders, and countries during project design, fieldwork, data collection, analyses, and writing. Most participants and coauthors are from Colombia, where the study took place, and researchers outside academia also participated in the project and publication.

