## Supplementary figures for "Flexibility in movement strategies of neotropical nectarivorous birds: insights from high-Andean hummingbirds and flowerpiercers"

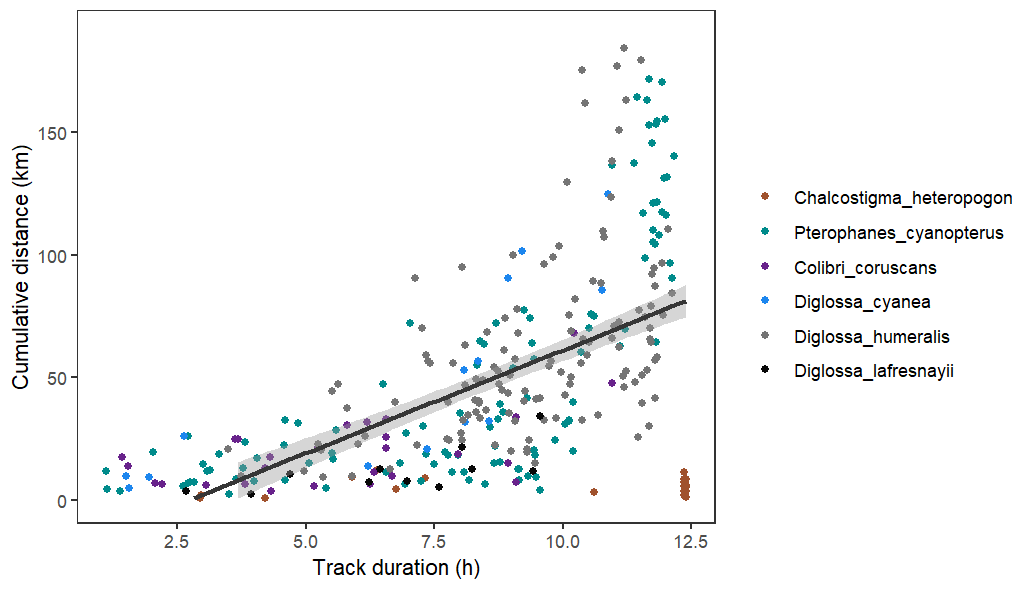

**Figure S1.** Correlation between daily cumulative distance and tracking duration. Colours of points show species according to legend and solid black line indicates the linear relationship between the two variables (r = 0.57 , p < 2.2e-16), with shading showing standard error.

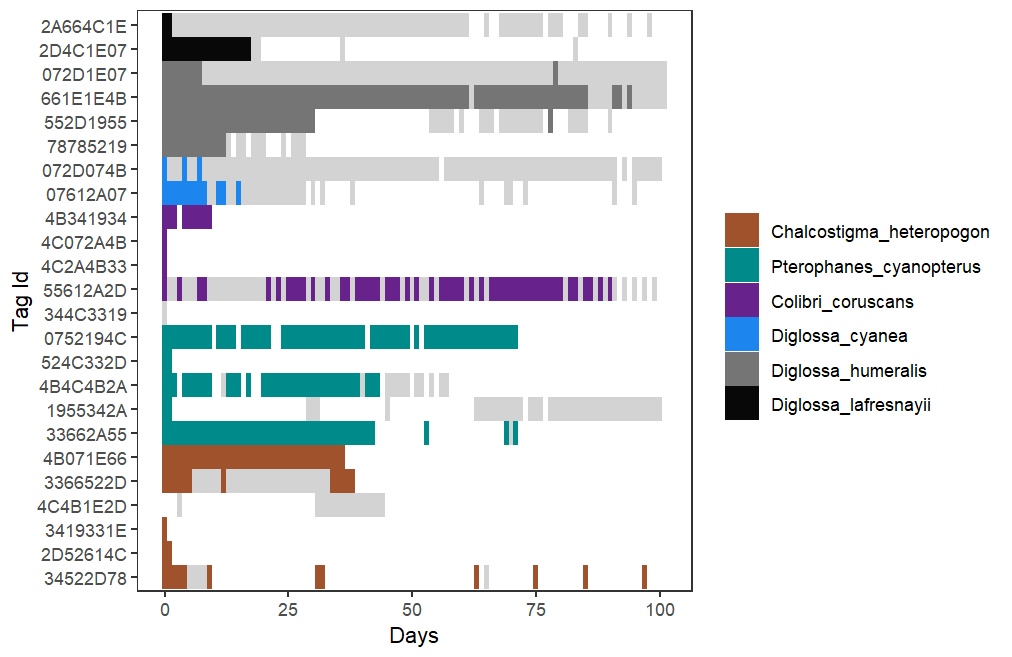

**Figure S2.** Duration of data received from 24 tagged birds. Coloured according to species when location estimates were possible, grey indicates radio signals were received but insufficient to estimate location (i.e. signals too low or detected by fewer than 3 nodes).

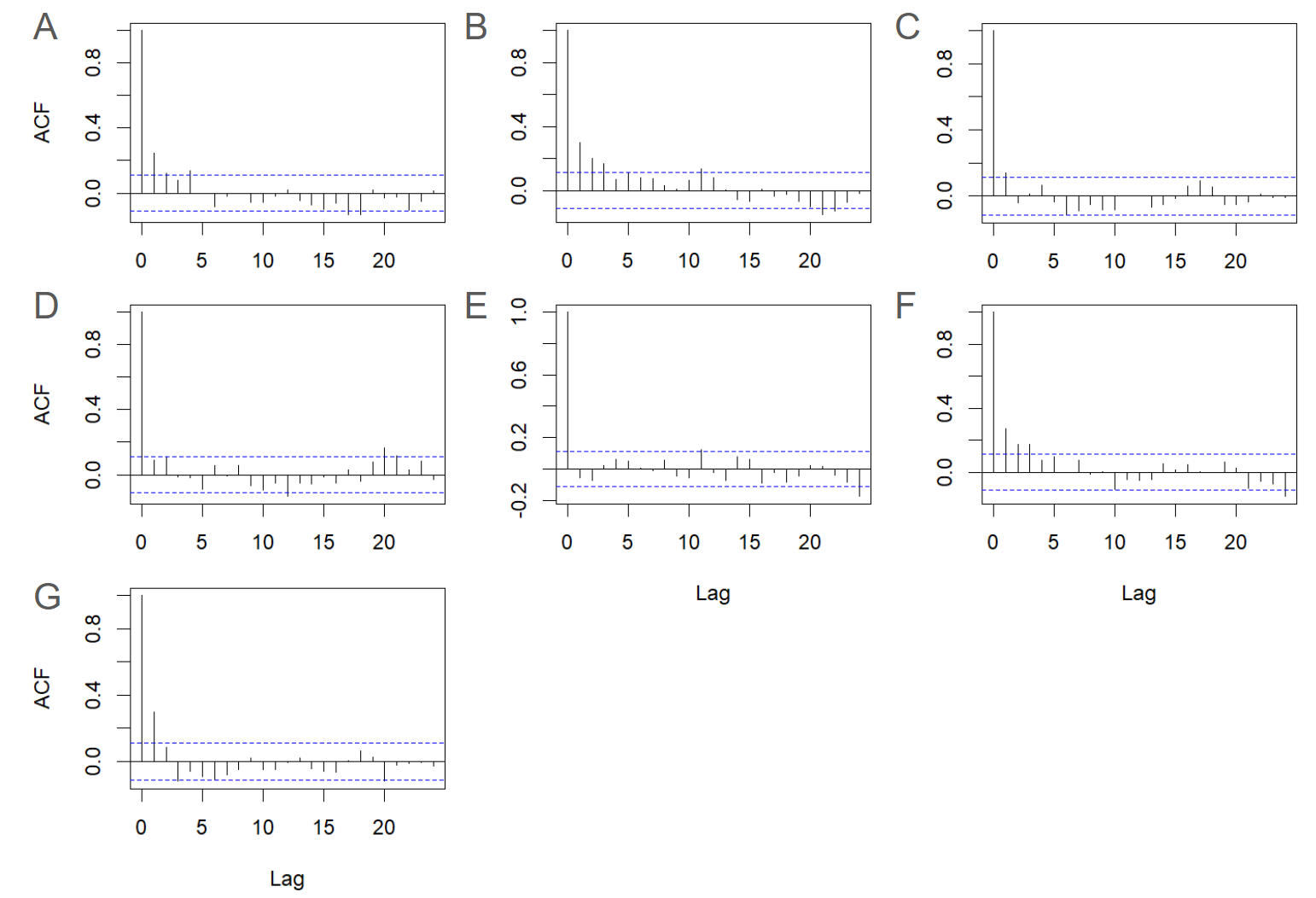

**Figure S3.** Autocorrelation plots of residuals in linear mixed effects models comparing seven movement metrics between hummingbirds and flowerpiercers. Panels refer to A) area, B) cumulative distance, C) centroid displacement, D) proportion of residence time, E) mean return time, F) revisitation rate, and G) proportion of forest within area.

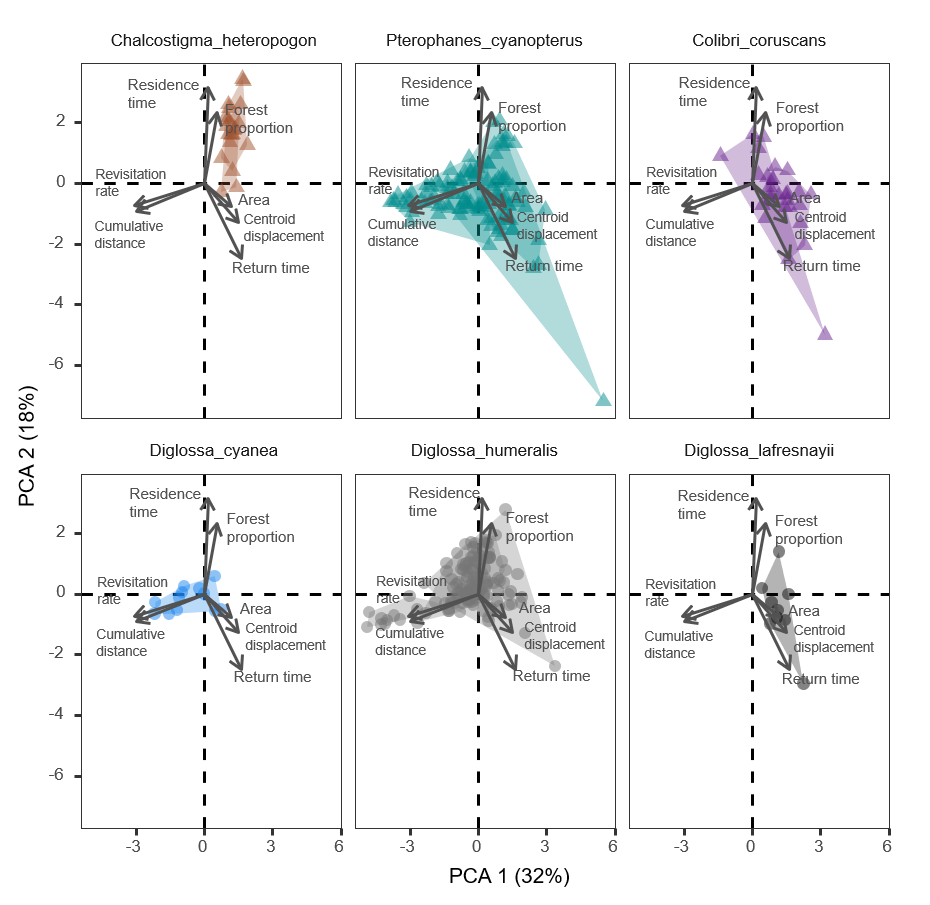

**Figure S4.** Principal components analyses (PCA) based on raw values by individuals, with values in parentheses indicating percent variance explained by each component. The first two components are shown, and points show daily values and are separated in different panels by species. The functional space occupied by an individual is shown as a convex hull. Note that variable loadings were multiplied by 5 to improve visualisation (raw values are available in Table SX).

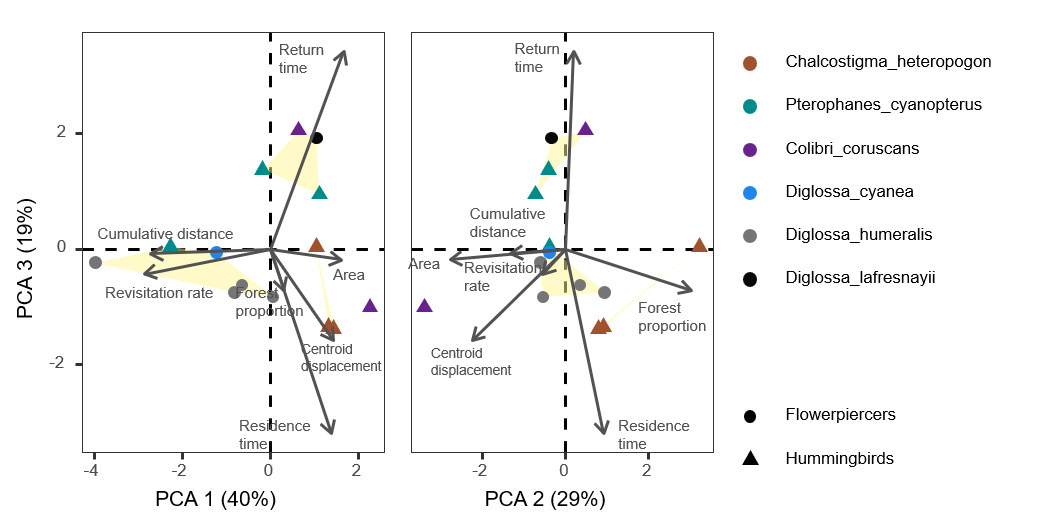

**Figure S5.** Principal components analyses (PCA) based on mean values by individuals, with values in parentheses indicating percent variance explained by each component, as in Figure 4A but adding a third component. Each point represents an individual, coloured according to species and bird family, as shown in the legend. Convex hulls in yellow show three main clusters identified. Note that variable loadings were multiplied by 5 to improve visualisation (raw values are available in Table SX).

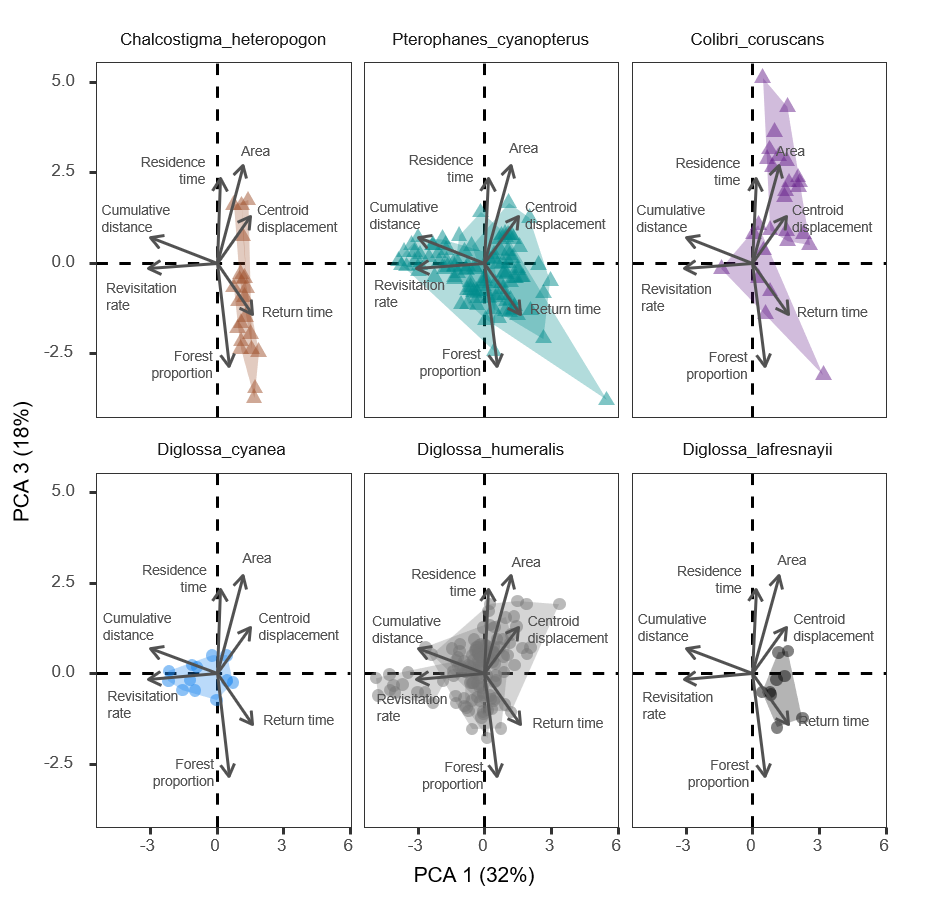

**Figure S6.** Principal components analyses (PCA) based on raw values by individuals, with values in parentheses indicating percent variance explained by each component. PC1 and PC3 are shown, and points show daily values and are separated in different panels by species. The functional space occupied by an individual is shown as a convex hull. Note that variable loadings were multiplied by 5 to improve visualisation (raw values are available in Table SX).

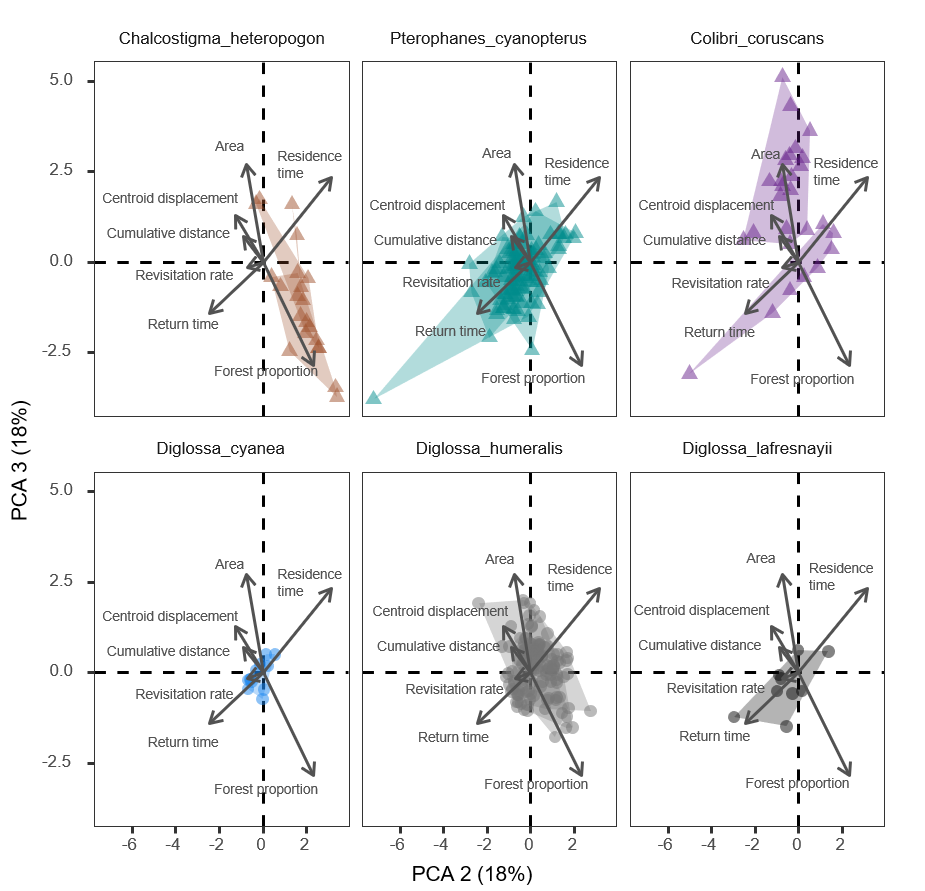

**Figure S7.** Principal components analyses (PCA) based on raw values by individuals, with values in parentheses indicating percent variance explained by each component. PC2 and PC3 are shown, and points show daily values and are separated in different panels by species. The functional space occupied by an individual is shown as a convex hull. Note that variable loadings were multiplied by 5 to improve visualisation (raw values are available in Table SX).

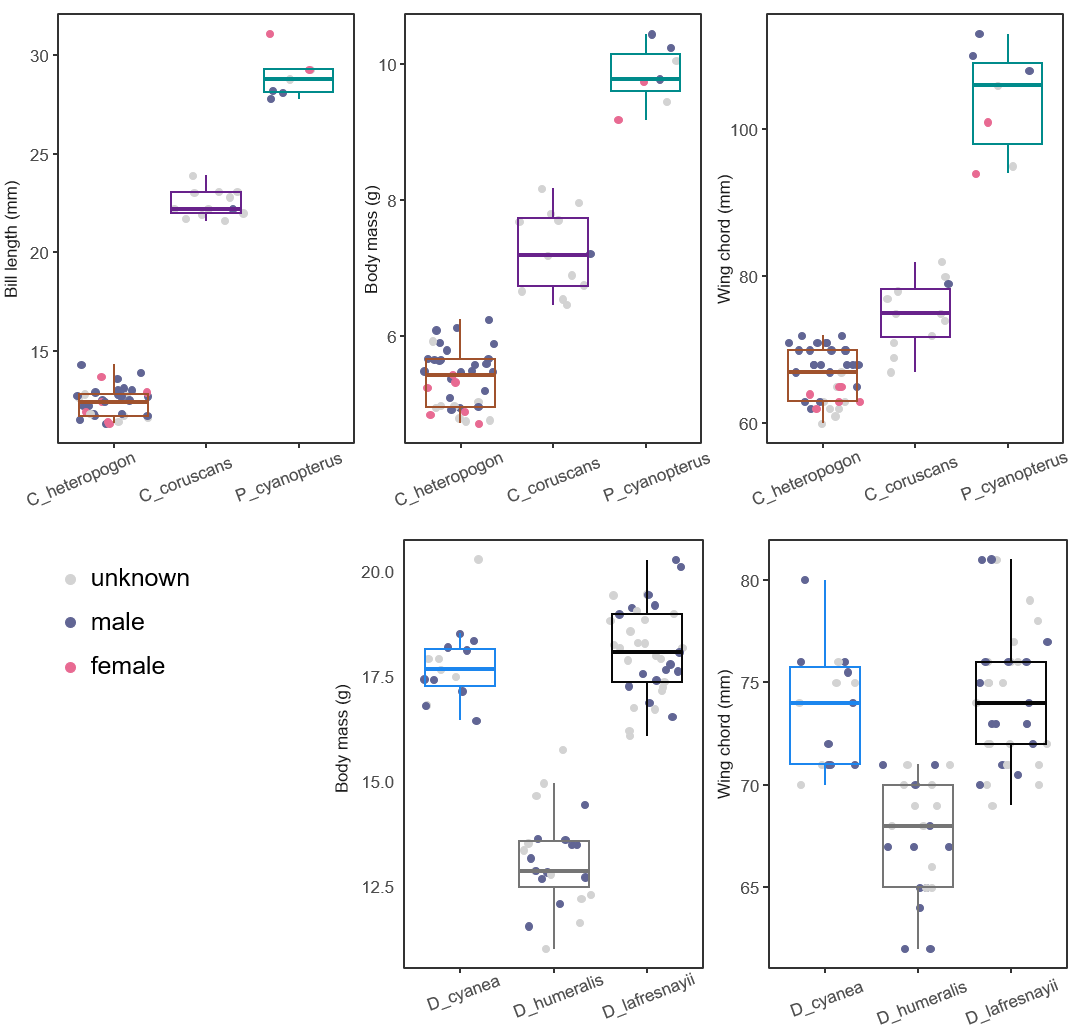

**Figure S8.** Body measurements of tagged species of hummingbirds and flowerpiercers, from mist netting effort between February 2022-August 2024. Points show individuals coloured by sex according to the legend, and boxplots indicate first and third quartiles with lower and upper box hinges, median values with middle horizontal lines, 1.5 interquartile ranges with vertical lines, and outliers with points.
